# A generalized synthetic control algorithm for sparse functional data

**DOI:** 10.64898/2026.02.23.707582

**Authors:** Lucy Shao, Kilian M. Pohl, Wesley K. Thompson

## Abstract

The Synthetic Control Method (SCM) and its interactive factor model generalizations (GSC) are powerful for estimating causal effects from panel data but are not easily applied when follow-up is irregular or sparse, common features of biomedical cohorts. We develop a Bayesian functional extension of GSC that treats each unit’s outcome path as a smooth latent trajectory and accommodates unequally spaced measurements. Trajectories are approximated using Functional Principal Components Analysis (FPCA), providing a data-driven basis that captures dominant patterns with minimal shape assumptions while borrowing strength across individuals. Within this representation, we learn unit and time latent factors jointly with FPCA scores from the control data, construct counterfactual trajectories for treated units, and quantify uncertainty via the posterior. Identification relies on a latent-factor/weak-trend condition and overlap of controls and treated units in the functional score space. Simulation studies varying donor pool and treated unit size and sampling density show that the proposed approach (a.k.a GSC-FPCA) yields low bias when sampling is irregular or sparse, with well-calibrated interval coverage across a broad range of scenarios. We apply the method to longitudinal neuroimaging data from the National Consortium on Alcohol and Neurodevelopment in Adolescence - Adulthood (NCANDA-A) study to estimate the effect of adolescent binge drinking on subsequent brain volumes. Leveraging from 1 to 9 observed time points per participant, GSC-FPCA produces stable counterfactuals and detects a negative impact on gray-matter volumes with sustained high levels of binge drinking. Our results demonstrate that embedding GSC within a functional framework enables robust causal inference in biomedical applications characterized by irregularly-spaced visits, limited observations, and complex outcome dynamics.

## 1 Introduction

The Synthetic Control Method (SCM) was developed to address the shortcomings of previous casual inference methods commonly applied to panel data [3]. One such method is Difference-in-Differences (DiD), which compares changes in outcomes over time between treated and control groups. (Note, throughout this article we use the term “treated” and “treatment” by convention; this can also refer to exposures, such as alcohol use, which are not generally considered treatments.) Specifically, DiD assumes outcomes follow a *two-way fixed effects* model:

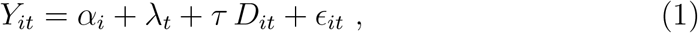

where *Y*_*it*_ is the outcome for unit *i* at time *t, α*_*i*_ are unit-specific fixed effects (capturing time-invariant differences across units), *λ*_*t*_ are time fixed effects (capturing time trends common to all units at time *t*), and *D*_*it*_ is a treatment indicator for whether unit *i* is exposed to treatment at time *t*. The error terms *ϵ*_*it*_ are assumed to be mean-independent of the treatment assignment process (no omitted time-varying confounders) [4]. Assuming also no interference between units and no anticipation of treatment, *τ* recovers the average treatment effect on the treated (ATT) [4]. In the simplest two-group, two-period setting, the DiD estimator of the ATT can be written as the difference in average outcome changes between the treated group *T* and control group *C*:

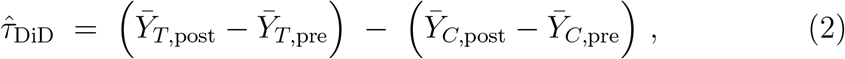

Where 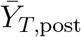 (respectively 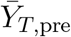) is the sample mean of the outcomes for the treated group in the post-treatment period, or *t* = 2 (respectively pretreatment period, or *t* = 1), and similarly for 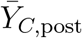 and 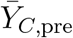 for the control group. The key identification assumption is that, in the absence of treatment, the treated units would have experienced the same outcome trend as the controls (parallel trends), which is modeled by *α*_*i*_ being time-invariant.

If there are time-varying unobserved factors that affect treated and control groups differently, this assumption fails and the DiD estimate will be biased. To relax this assumption, the Synthetic Control Method (SCM)[3, 1, 2] constructs a weighted combination of control units (a “synthetic” control) that closely reproduces the treated unit’s outcome trajectory in the pretreatment period. By matching the treated unit’s pre-treatment outcomes (and possibly covariates) as closely as possible, the synthetic control is meant to approximate what the treated unit’s outcome path would have been in the absence of the intervention. This approach allows for unobserved confounders to have time-varying effects on outcomes.

In the simplest case, suppose there is one treated unit (say unit *i* = 1) and *n*−1 potential control (donor) units (*i* = 2, …, *n*) that remain untreated throughout. Let *T*_0_ be the last time period before the treatment begins. SCM seeks weights *w*_2_, …, *w*_*n*_ for the control units such that the weighted control outcomes closely track the treated unit’s outcomes for *t* ≤ *T*_0_. For example, one approach is to solve:

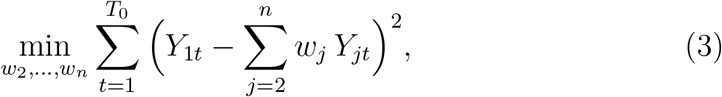

subject to *w*_*j*_ ≥ 0 and 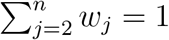. These constraints restrict the synthetic control to be a convex combination of actual control units, which avoids extrapolation beyond the support of the observed data. The optimization uses only pre-intervention data. After determining the weights *w*_*j*_, the counter-factual outcome for the treated unit in the post-treatment period *t > T*_0_ is estimated as the synthetic control’s outcome (a weighted sum of the donors). The treatment effect for unit 1 at time *t* is then:

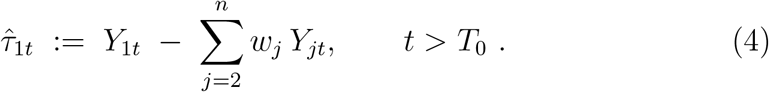

By construction, the synthetic control ∑_*j*_ *w*_*j*_*Y*_*jt*_ mimics *Y*_1*t*_ for *t* ≤ *T*_0_. Under the assumption that this good pre-treatment fit implies the synthetic control captures the effects of the relevant confounding factors affecting unit 1 in the post-treatment period, the difference 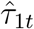 provides an approximately unbiased estimate of the treatment effect for *t* > *T*_0_.

SCM has proven useful in settings with a single (or very few) treated units and many available control units, especially for aggregate-level case studies [1, 2]. In practice, SCM requires a sufficiently long pre-intervention period and a close pre-treatment fit to unbiasedly estimate causal effects. It may have larger bias and higher variance if only a few pre-treatment periods are available or if the treated unit’s pre-treatment trajectory cannot be well reproduced by any convex combination of controls [1]. In those cases, the synthetic control may not adequately approximate the counterfactual.

One way to interpret SCM is via latent factor models for the outcome. For instance, Abadie et al. [1] consider a model for the untreated potential outcomes:

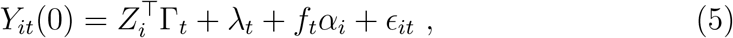

where *Z*_*i*_ is a vector of observed covariates (with time-varying coefficients Γ_*t*_) and *f*_*t*_*α*_*i*_ represents unobserved factors (with *α*_*i*_ an unobserved factor loading for unit *i* and *f*_*t*_ a time-varying common factor). This latent factor structure generalizes the DiD model: if we forced *f*_*t*_ = *f* to be constant over *t*, then *fα*_*i*_ is just a unit fixed effect and we are back to parallel trends. By allowing *f*_*t*_ to vary with *t*, unobserved confounders can have time-varying impacts. Intuitively, the synthetic control weights are chosen such that the weighted controls replicate the treated unit’s combination of these latent factors in the pre-treatment period. Under standard conditions (the latent factor model holds and the pre-treatment fit is good), the synthetic control provides an approximately unbiased estimate of the treated unit’s counterfactual path [4].

While SCM constructs a nonparametric counterfactual for one treated unit at a time, *generalized* synthetic control approaches take a more model-based perspective by explicitly estimating a latent factor structure for the panel data. Building on the econometric literature of interactive fixed effects [e.g., 6], these methods assume the outcome can be decomposed into observed covariates and a low-dimensional set of latent factors that drive common trends. A general model for the no-treatment outcome is:

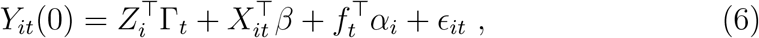

where *X*_*it*_ are observed time-varying covariates (with coefficient vector *β*), and *α*_*i*_ and *f*_*t*_ are *F*-dimensional vectors of latent factor loadings and common factors, respectively. If *F* = 1 and *f*_*t*_ is constant over time, then *f*_*t*_*α*_*i*_ reduces to a unit fixed effect, recovering the classical DiD setup as a special case. (The time trend *λ*_*t*_ is absorbed into the first term on the right side of the equation, if *Z*_*i*_ includes an intercept.) With *F >* 1 and time-varying *f*_*t*_, this model can capture rich temporal dynamics.

The Generalized Synthetic Control (GSC) method proposed by Xu [19] leverages such factor models for causal inference. In essence, GSC uses the entire panel of control units and time periods to learn the latent factors *f*_*t*_ and factor loadings *α*_*i*_. For example, one can estimate *f*_*t*_ and *α*_*i*_ by applying principal components analysis to the matrix of control outcomes [6]. Once the common factors are estimated from the control group, a treated unit’s factor loadings can be inferred from its pre-treatment outcomes. These estimated loadings are then used to predict the treated unit’s counterfactual outcomes for *t > T*_0_. By explicitly modeling the latent structure (*α*_*i*_, *f*_*t*_), this approach automatically balances the treated unit’s trajectory with the controls in a way analogous to optimizing weights in SCM, but within a parametric framework. Related work by Athey et al. [5] cast the counterfactual prediction problem as one of *matrix completion*. By assuming the complete panel (with missing entries for treated outcomes post-treatment) has low rank (a form of factor model), they applied convex optimization to fill in the missing entries. This matrix completion estimator is closely related to GSC but uses a different estimation strategy. Simulation studies have shown that when the latent factor model holds, these approaches (GSC, matrix completion) can outperform basic SCM in terms of bias and mean squared error.

The advantages of the GSC framework include the ability to handle multiple treated units and staggered treatment adoption in a unified way. Unlike the classic SCM, which is often used for a single treated unit, factor-model-based methods can accommodate many treated units (possibly with treatments beginning at different times) by incorporating treatment indicators into the model or allowing *α*_*i*_, *f*_*t*_ to capture heterogeneous exposure patterns. By pooling information across all units and time points to estimate the latent structure, GSC can achieve greater statistical efficiency than analyzing one treated case at a time. Moreover, the factor model approach provides a path to standard inferential techniques: under large *N* asymptotics, one can derive standard errors for the estimated factors, loadings, and treatment effects [e.g., 6, 19]. This is in contrast to the original SCM literature, which often relied on placebo tests and other non-parametric inference approaches (e.g., permutation of the treatment assignment) due to having only a single treated unit [1].

In biomedical and other observational studies, data are frequently collected at irregular time intervals or subject-specific schedules. Standard panel methods require aligning everyone to common time points or aggregating to fixed periods, potentially discarding information. Here, we extend the GSC approach to accommodate irregularly sampled longitudinal data by leveraging flexible trajectory models from functional data analysis. In particular, we propose to use Functional Principal Component Analysis (FPCA) to model trajectories in the sparse functional data setting. FPCA provides a data-driven basis that can capture underlying trajectory patterns with minimal assumptions on shape [20, 11, 18]. This allows us to relax strict parametric assumptions about outcome dynamics while still borrowing strength across individuals. In Section 2, we present our novel extension of GSC for irregularly observed functional data using FPCA. Section 3 describes simulation experiments that benchmark our new method across a range of data-generating scenarios (varying latent factor strengths, donor pool sizes, and pre-treatment lengths). In Section 4, we apply the methods to an empirical case study examining the impact of adolescent binge drinking on later brain volume, comparing estimates and inference from each approach, using data from the National Consortium on Alcohol and Neurodevelopment in Adolescence - Adulthood (NCANDA-A) study [7]. Finally, Section 5 discusses the limitations of our method, possible extensions (e.g., incorporating nonlinear effects or multiple outcome trajectories), and practical guidance for researchers.

## 2 Method

### 2.1 Functional Generalized Synthetic Control Model

We propose a continuous-time generalized synthetic control method suitable for sparse functional (longitudinal) data [13, 17]. Consider a study with a treated (exposed) group 𝒯 and a control (unexposed) group 𝒞. Let *N*_tr_ = |𝒯| be the number of exposed individuals and *N*_co_ = |𝒞| be the number of controls, with total sample size *N* = *N*_tr_ + *N*_co_. For each subject *i*, let *Y*_*i*_(*t*) denote the outcome of interest at time *t*. Subjects may be observed at an individual-specific set of time points 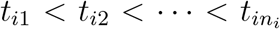, which can be irregularly spaced. We assume each exposed subject *i* ∈ 𝒯 has a well-defined treatment start time *T*_0,*i*_, such that *Y*_*i*_(*t*) for *t < T*_0,*i*_ represents the pre-exposure trajectory and *Y*_*i*_(*t*) for *t* ≥ *T*_0,*i*_ is the post-exposure trajectory under treatment (i.e., once exposed always exposed). Control units are never exposed during the study period (conceptually *T*_0,*i*_ = ∞ for *i* ∈ 𝒞).

We adopt a functional data model for each individual’s trajectory [11]. Let *µ*(*t*) be the overall mean function of the outcome, and let *f*:= (*f*_1_(*t*), *f*_2_(*t*), …, *f*_*k*_(*t*)) be a vector consisting of the first *k* functional principal component (FPC) functions capturing the major modes of variation around *µ*(*t*) [11, 18]. For identifiability, the *k* FPC basis functions are orthonormal, i.e., ∫ *f*_*j*_(*t*)*f*_*ℓ*_(*t*) *dt* = *δ*_*jℓ*_, where *δ*_*jℓ*_ is the Kronecker delta function. We assume an expansion of each individual’s trajectory in terms of these functions. In addition to the latent trajectory components, we allow observed covariates: let *X*_*i*_(*t*) be a vector of time-varying covariates with linear effects *β* on *Y*_*i*_ and let *Z*_*i*_ be a vector of time-invariant covariates for subject *i* with time-varying coefficient functions Γ(*t*). Finally, let *D*_*i*_(*t*) be a treatment indicator function (equal to 1 if subject *i* is exposed to treatment by time *t*, and 0 otherwise). We allow a treatment effect *δ*(*t*^′^) that can vary flexibly with time since exposure, denoted by *t*^′^ = *t* − *T*_0_. While we could consider exposure effects that depends more generally on time of exposure (i.e., *δ*(*t, T*_0_)), for simplicity this model assumes that only time since is exposure is relevant for effect sizes.

Formally, we assume the following model for the observed outcome trajectory of individual *i*:

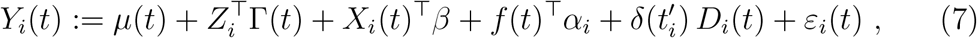

for *t* in the observation interval (e.g. *t* ∈ [0, *T*_max_]). The vector *α*_*i*_:= (*α*_*i*1_, …, *α*_*ik*_)^⊤^ consists of the subject-specific scores (random coefficients) on the FPCs for individual *i* [11], which we assume has a mean zero and a diagonal covariance matrix. The term *ε*_*i*_(*t*) represents a residual measurement error process, assumed to be mean zero and uncorrelated across subjects (and uncorrelated with the covariates and latent factors) with variance *σ*^2^ for all *t*. The function 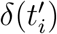 represents the treatment effect as a function of time since exposure 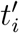 for the *i*th participant. For *t < T*_0,*i*_ (before treatment), *D*_*i*_(*t*) = 0 so 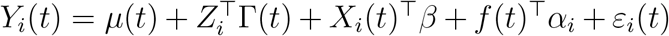. For control units (who serve as the donor pool for constructing counterfactuals), *D*_*i*_(*t*) ≡ 0 for all *t*, so their outcomes are always given by the no-treatment trajectory.

To implement this model, we represent the mean function *µ*(*t*), the time-varying coefficients Γ(*t*) and the functional components *f* (*t*) in a basis expansion. Let *b*(*t*) = (*b*_1_(*t*), …, *b*_*q*_(*t*))^⊤^ be a chosen spline basis (or other suitable basis) of dimension *q* for functions on the time domain. We will approximate *µ*(*t*), each component of Γ(*t*), and each FPC *f*_*j*_(*t*) in this basis. For subject *i* with observation times 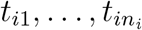, define *B*_*i*_ as the *n*_*i*_ × *q* matrix of basis function evaluations at those times (i.e., the (*r, ℓ*) element of *B*_*i*_ is *b*_*ℓ*_(*t*_*ir*_)). We choose the basis *b*(·) such that, on a fine grid covering the domain, the matrix of basis evaluations *B* satisfies *B*^⊤^*B* ≈ *I*_*q*_ (approximately orthonormal basis functions). In practice, one can use an orthonormalized spline basis.

Using this basis expansion, we can rewrite Eq. (7) in vector/matrix form for the observations of subject *i*. Let 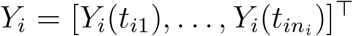 be the out-come vector for subject *i*. Likewise, define the vector of treatment indicators 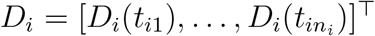 (this will be a vector of 0’s for a control or preexposure period, switching to 1’s after exposure for a exposed unit). Let 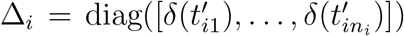 be the *n*_*i*_ × *n*_*i*_ diagonal matrix of treatment effects evaluated at the observed times since exposure onset for the *i*th unit. Define 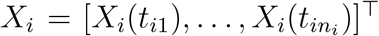 as the *n*_*i*_ × *p* matrix of observed values of the time-varying covariates for subject *i*, with corresponding *p*-vector of unknown coefficients *β* = (*β*_1_, …, *β*_*p*_)^⊤^. Also define *Z*_*i*_ = (*Z*_*i*1_, …, *Z*_*ir*_)^⊤^ as the *r*-dimensional vector of time-invariant covariates for subject *i*. Then Eq. (7) evaluated at the observed time points can be written as:

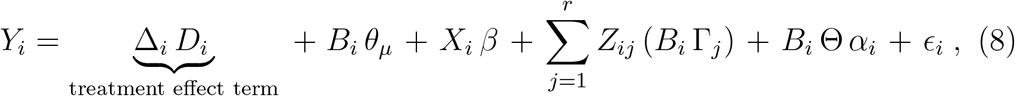

where *θ*_*µ*_ is the *q*×1 coefficient vector for the mean function expansion (*µ*(*t*) ≈ *b*(*t*)^⊤^*θ*_*µ*_), Γ_*j*_ is the *q* × 1 coefficient vector for the *j*th covariate effect function (Γ_*j*_(*t*) ≈ *b*(*t*)^⊤^Γ_*j*_), and Θ is the *q* × *k* matrix whose columns are the spline coefficient vectors for the FPC functions 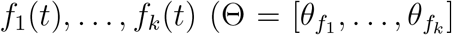,so 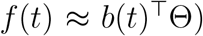. For a exposed unit *i, D*_*i*_ selects out the post-treatment entries and multiplies by the effect function values, whereas for a control *i* or pre-treatment period, *D*_*i*_ is zero and Δ_*i*_ *D*_*i*_ contributes nothing. We impose the identifiability constraint Θ^⊤^Θ = *I*_*k*_ (so that the FPCs are orthonormal in the chosen basis representation). The primary estimand of interest is the average treatment effect on the treated (ATT) as a function of time since exposure *t*^′^ ≥ 0, define, i.e.,

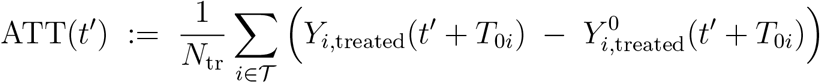

### 2.2 Estimation strategy

We now outline how we estimate the model parameters and infer the treatment effect using a Bayesian approach with a Gibbs sampler. The goal is to use the control individuals (and pre-treatment data from treated individuals) to learn the regression parameters *β* and functional components *µ*(*t*), Γ(*t*), *f*_*j*_(*t*) and variance parameters, then to predict each treated individual’s counterfactual trajectory 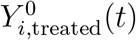 for *t* ≥ *T*_0_. The treatment effect for treated unit *i* at time *t*^′^ since treatment onset *T*_0,*i*_ is then estimated as

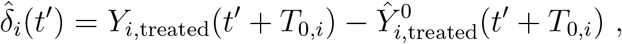

and the ATT at time *t* is the average of 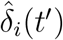 over *i* ∈ 𝒯.

### 2.3 Bayesian FPCA model for the counterfactual trajectory

For estimation, we consider the following generative model for the counter-factual outcomes (setting *D*_*i*_(*t*) = 0) with *p* time-varying predictors *X*_*ij*_(*t*) and *r* time-invariant predictors *Z*_*ij*_:

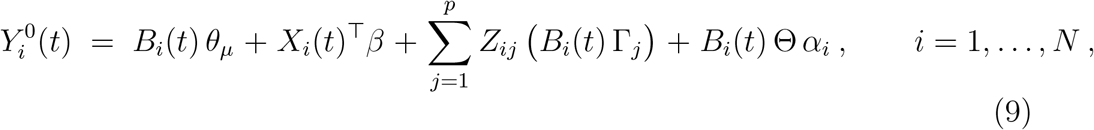

with the orthonormality constraint Θ^*T*^ Θ = *I*_*k*_ as before. Here 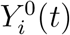 represents the latent no-treatment outcome for subject *i* at time *t* (for controls this is just the smoothed *Y*_*i*_(*t*), for treated units this corresponds to their noise-free outcome if treatment had never occurred). We place the following prior distributions on the model parameters:

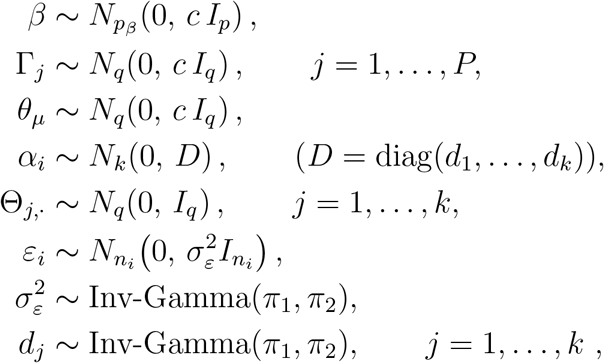

where *N*_*m*_(0, Σ) denotes an *m*-dimensional multivariate normal with mean 0 and covariance Σ. In words, we assume relatively diffuse Gaussian priors (with variance *c*) for the fixed-effects coefficients *β*, Γ_*j*_, *θ*_*µ*_, and independent inverse-gamma priors for the residual variance 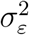 and each FPC score variance *d*_*j*_. The prior on the columns Θ_*j*,·_ of the FPC basis (in the spline space) is taken as standard normal *N* (0, *I*_*q*_), subject to the orthonormality constraint Θ^⊤^Θ = *I*_*k*_ (enforced in the sampling scheme). We fit the latent functional components (*µ*(*t*), *f*_*j*_(*t*), etc.) using only the control units’ data. The number of retained FPCs is determined by fitting models with *k* = 1, …, *k*^′^ (with *k*^′^ ≤ *q*) and computing the PSIS–LOO estimate of expected log predictive density, elpd_loo_, reported as the LOO information criterion LOOIC = −2 elpd_loo_, for each *k* [16]. We retain the *k* with the lowest LOOIC (equivalently, highest elpd_loo_), unless differences are within 1–2 standard errors, in which case the simpler model is preferred.

Details of the Gibbs sampling algorithm are given in the Supplementary Materials section.

## 3 Monte Carlo Simulations

We conducted simulations under various scenarios to evaluate the finite sample performance of the proposed method. We generated longitudinal data for *N* = 50, 100, and 200 subjects (with *N*_tr_ = 20, 40, 80 treated). The observation times for each subject were randomly drawn in the interval [0, 1]. We considered three levels of sparsity in observation times: in scenario A, each subject’s number of observations *n*_*i*_ was ∼ Poisson(2) (with a minimum of 2 observations per subject); in scenario B, *n*_*i*_ ∼ Poisson(4) (min 3); in scenario C, *n*_*i*_ ∼ Poisson(7) (min 4). The maximum potential number of observations per subject was capped at *T*_max_ = 10, but most subjects had far fewer time points in these sparse scenarios.

We generated the latent trajectory components as follows. The underlying mean function *µ*(*t*) was taken to be a smooth nonlinear curve (see Figure 1 for a visualization in one scenario). We used *q* = 3 cubic spline basis functions for *B*_*i*_ in these simulations. We set *k* = 2 principal components, with the first FPC *f*_1_(*t*) proportional to cos(*t*) and the second *f*_2_(*t*) proportional to sin(*t*) over *t* ∈ [0, 2*π*] (normalized to unit *L*^2^ norm on [0, 2*π*]; see Figure 2). The associated subject scores *α*_*i*1_ were generated from 𝒩 (0, 1) and *α*_*i*2_ from 𝒩 (0, 4), so the second component had larger variability. The measurement error standard deviation was *σ*_*ε*_ = 1.0. For covariates, we included one binary time-invariant covariate *Z*_1*i*_ ∼ Bernoulli(0.5) (along with an intercept term implicitly included in *µ*(*t*)) and two time-varying covariates: *X*_*i*1_(*t*) drawn i.i.d. 𝒩 (0, 1) and *X*_*i*2_(*t*) drawn i.i.d. Bernoulli(0.5) at each observed time. All regression coefficients were set to 1. Thus, the true data-generating model for the no-treatment outcomes was:

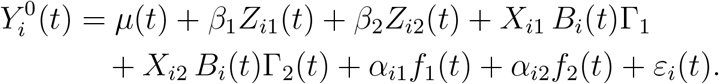

with *β*_1_ = 1, *β*_2_ = 1, Γ_1_ ≡ 1, and Γ_2_ ≡ 1. Figure 1 shows an example of the population mean function *µ*(*t*) used (for *N* = 100, scenario B), and Figure 2 shows the two FPC curves. 3 visualizes the average treatment effect(for *N* = 100, scenario B) across simulation runs grouped by time rounded to the nearest 0.1.

**Figure 1.**
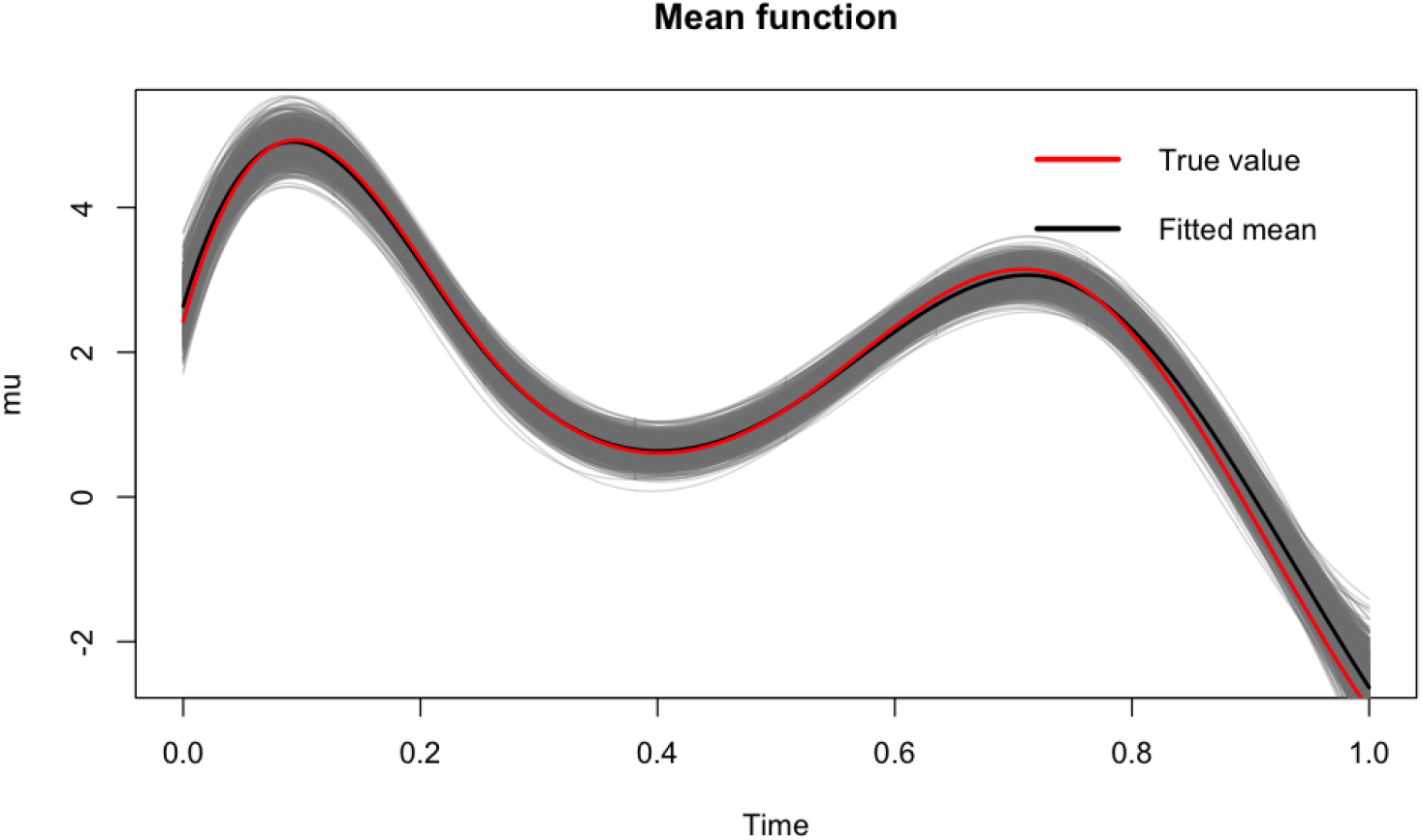
True mean function *µ*(*t*) (N=100; average of 4 time points per subject). The red curve denotes the true mean, the grey curves denote fitted means from individual simulation runs, and the black curve denotes the fitted mean averaged across simulation runs.

**Figure 2.**
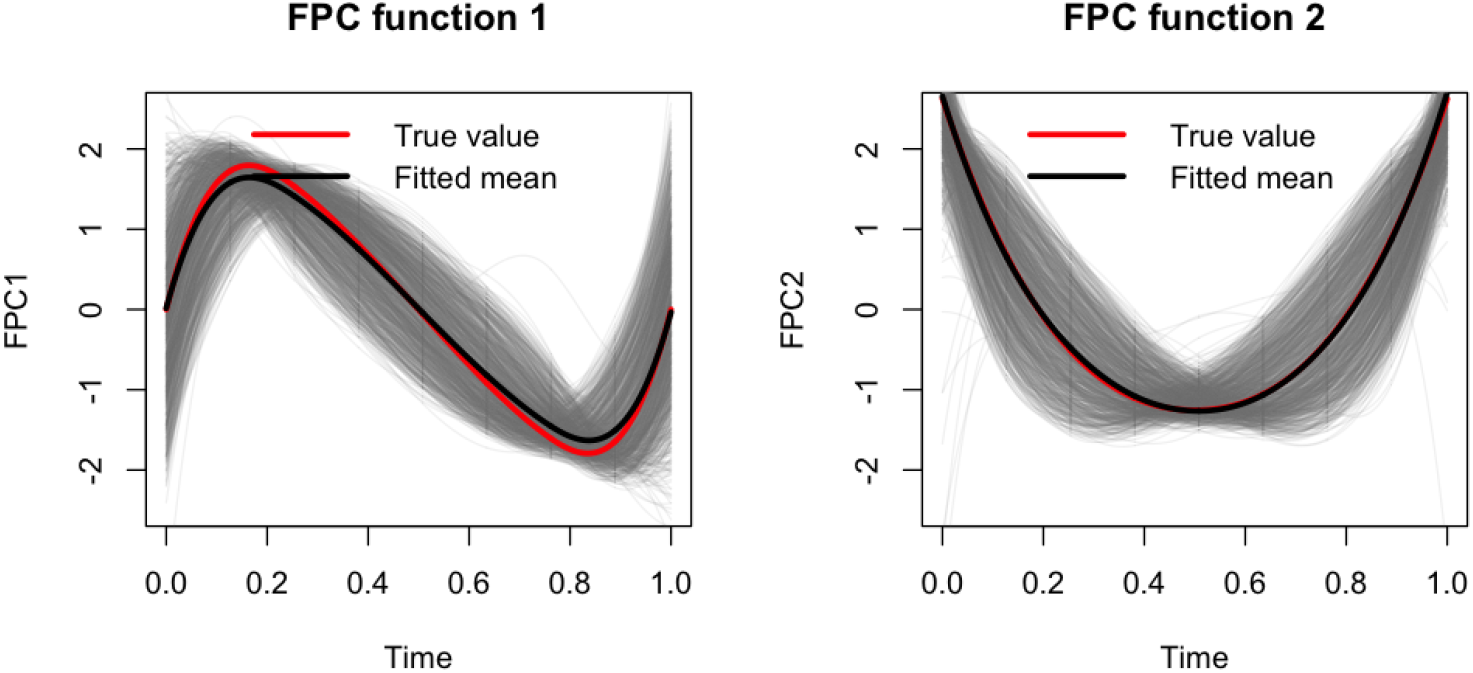
True first two functional principal component curves (*f*_1_(*t*) and *f*_2_(*t*)) fitted in the simulations (N=100; average of 4 time points per subject). The red curve denotes the true value, the grey curves denote fitted means from individual simulation runs, and the black curve denotes the fitted value averaged across simulation runs.

**Figure 3.**
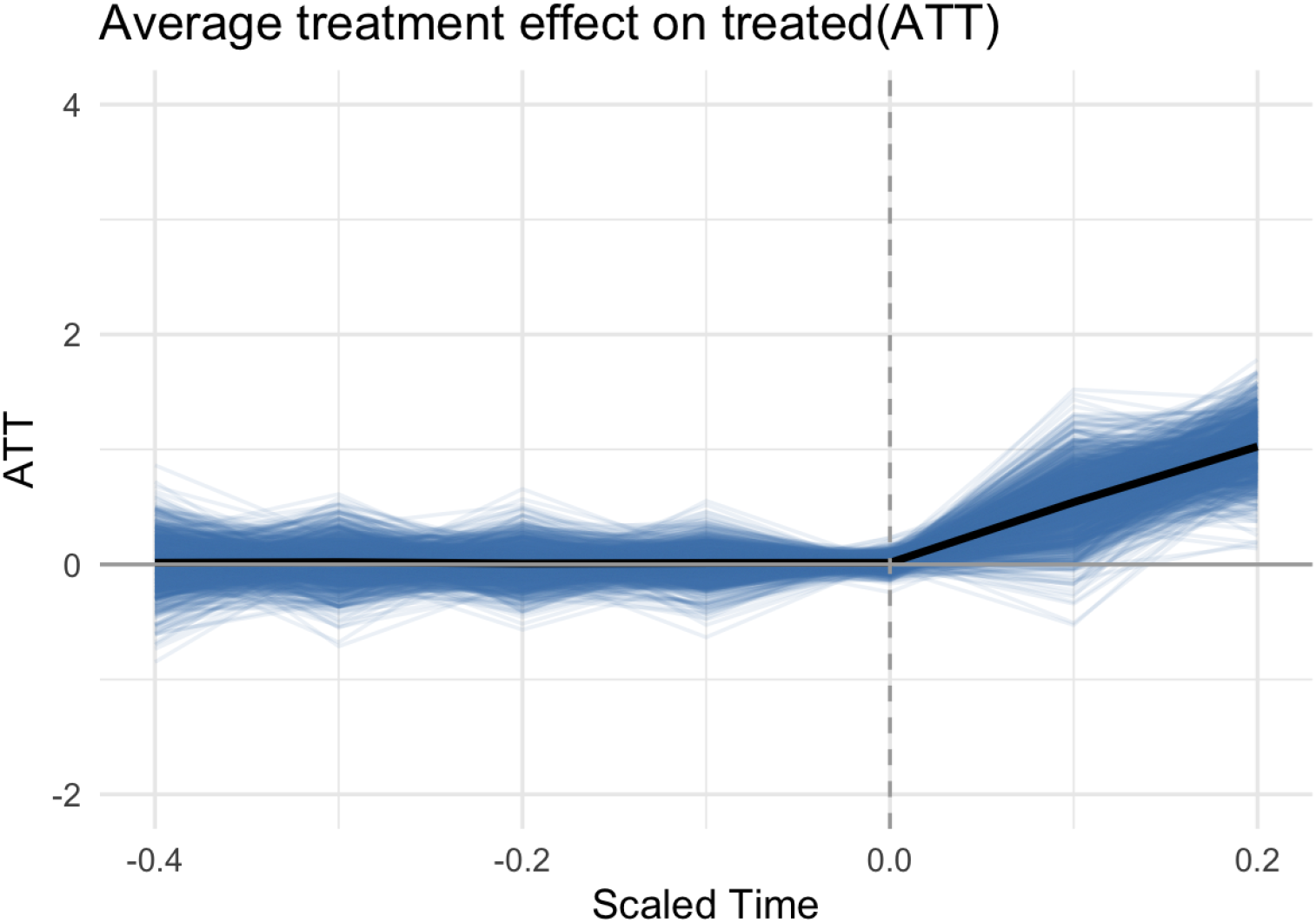
Average treatment effect grouped by every 0.1 nearest scaled time after treatment started. (*N*_*control*_ = 100, average 4 time points)).

We introduced a treatment effect for the treated units. We assumed treatment begins at time *T*_*i*,0_ = ⌊2*T*_*i*,max_*/*3⌋ (approximately two-thirds through the observation period). The true treatment effect for a treated unit *i* at time *t* ≥ *T*_*i*,0_ was specified as a function of time since exposure: *δ*_*i*_(*t*) = *G*(*t*− *T*_*i*,0_) with *G*(Δ) = 5 × Δ. In other words, the causal effect causes a linear increase (or decrease) in the outcome of 5 units per unit time after exposure. This deterministic effect is the same for all treated individuals in our simulations for simplicity. The observed outcome for a treated unit thus equals the generated 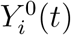 plus this treatment effect after *T*_0_.

We applied our method to the simulated datasets to estimate the ATT over time. We summarize results focusing on the estimation of the ATT shortly after treatment initiation. The true ATT at a specific time *t* (with *t* = *T*_0_ + Δ) is *G*(Δ) by construction (e.g., at Δ = 0.1 after exposure, ATT = 5 × 0.1 = 0.5). We evaluated the bias and mean squared error (MSE) of the estimated ATT at the treatment time for each subject over 1000 Monte Carlo repetitions for each scenario. We also computed coverage of 95% posterior credible intervals (or confidence intervals) for ATT.

Table 1 reports the mean bias, MSE, and coverage for the ATT estimate at the at time 0.1(rounded) after treatment starts after treatments starts under various scenarios. Each entry is based on 1000 simulated datasets. We vary the total sample size and sparsity of observations as described (scenario A: very sparse, B: moderate, C: relatively dense on average).

**Table 1:** Simulation results for ATT at time 0.1(rounded) after treatment starts. *N*_tr_ is the number of treated; *N*_co_ is the donor pool size. Bias and MSE are reported as mean (SD); empirical coverage is for a nominal 95% interval.

| $N_{\text{tr}}$ | $N_{\text{co}}$ | Min/Avg $n_i$ | Bias mean (SD) | MSE mean (SD) | 95% Cov. ($N_{\text{tr}}=100$) |
| --- | --- | --- | --- | --- | --- |
| 20 | 50 | 2 / 2 | 0.136 (0.830) | 2.536 (1.808) | 0.918 |
| 20 | 50 | 3 / 4 | 0.018 (0.697) | 1.957 (1.399) | 0.930 |
| 20 | 50 | 4 / 7 | 0.001 (0.469) | 1.458 (0.812) | 0.946 |
| 40 | 100 | 2 / 2 | 0.010 (0.644) | 2.167 (1.321) | 0.920 |
| 40 | 100 | 3 / 4 | 0.038 (0.493) | 1.803 (0.982) | 0.938 |
| 40 | 100 | 4 / 7 | 0.009 (0.321) | 1.410 (0.543) | 0.950 |
| 80 | 200 | 2 / 2 | 0.034 (0.438) | 2.048 (0.884) | 0.929 |
| 80 | 200 | 3 / 4 | 0.007 (0.346) | 1.757 (0.649) | 0.934 |
| 80 | 200 | 4 / 7 | -0.012 (0.231) | 1.396 (0.371) | 0.939 |

The results indicate that our method produces approximately unbiased ATT estimates across scenarios, with bias on the order of a few hundredths (relative to true ATT of 0.5 at *T*_0_ + 0.1). As expected, estimation accuracy improves as the sample size and the density of observations increase. For example, in the most sparse and smallest scenario (*N*_co_ = 50, avg. 2 points per curve), the ATT estimator has an MSE around 2.7; this drops to about 1.4 in the largest, densest scenario (*N*_co_ = 200, avg. 7 points). The empirical coverage generated from 100 treated units of 95% intervals are mostly close to nominal in all cases (around 93–95%), except for smaller sample size and most sparse case(0.918, 0.920), suggesting that the Bayesian posterior intervals are well calibrated. We also observed that bias and MSE decrease as the control pool size *N*_co_ grows, highlighting the benefit of borrowing information from more donors. Furthermore, having slightly more measurements per subject (scenario C vs A) substantially improves performance, reflecting the advantage of even moderately dense functional data for learning individual trajectories.

## 4 Empirical Example

To demonstrate the GSC-FPCA method on a real data example, we apply our method to examine the impact of adolescent binge drinking on subsequent brain volumes. Data were obtained from the National Consortium of Alcohol on NeuroDevelopment in Adolescence to Adulthood (NCANDA-A; NDAR Data Collection ID 4513; Release: NCANDA NIAAADA 09Y IMAGING V01). NCANDA began in 2012 when five United States research sites (University of Pittsburgh, Stanford Research International, Duke University, Oregon Health and Science University, and the University of California, San Diego) enrolled 831 individuals to participate in a baseline visit and nine annual follow-ups. The study used an accelerated longitudinal cohort design with three age bands (12.0-14.9, 15.0-18.9, and 18.0-21.9 years) to capture neurodevelopment from adolescence to early adulthood. Most participants had no or low drinking experience at baseline (*N* = 692), while the remainder reported a drinking history that exceeded study thresholds for their age cohort [7]. We limited the age range of the analyzes in this longitudinal study from 12 to 21 years of age. We focus on *N* = 628 members without /low drinking at the beginning of the study with age ranges of 12 to 21 years. Each site obtained IRB approval; minors provided written assent with parental consent, and adults provided written informed consent at each annual visit.

The Customary Drinking and Drug Use Record (CDDR) [8] was collected annually to characterize current and past alcohol/substance use, capturing drinking frequency, average/maximum drinks, and binge drinking frequency. A binge drinking episode is defined as consuming 4/5 drinks (female/male) on one occasion (NIAAA). Participants reporting any binge drinking at baseline were excluded from the current analyses. Binge drinking episodes were assessed as the number of binge drinking episodes since the last assessment.

Magnetic Resonance Imaging (MRI) data were obtained from NCANDA-A participants. All sites followed standardized high-resolution structural MRI protocols using 3 T scanners. Three sites (SRI, Duke, and UCSD) employed GE MR750 scanners with 8-channel head coils, while two sites (UPMC and OHSU) used Siemens TIM-Trio scanners with 12-channel head coils. To ensure scanner consistency and monitor temporal variation, each site acquired an ADNI phantom at regular intervals [12], facilitating inter-site harmonization. T1- and T2-weighted images were acquired in the sagittal plane and processed via the SIBIS pipeline that included bias-field correction and skull stripping [12]. Intracranial volume (ICV) was defined based on the SRI24 atlas [14]. From these scans, 42 bilateral Desikan–Killiany regions of interest (ROIs) were derived for analysis, including 34 gray-matter and 8 white-matter ROIs, using Freesurfer [9].

Our exposure of interest was having at least 12 binge drinking episodes in the past year, i.e., averaging one or more such episodes a month. We kept as our exposed group all individuals who at some point during the study met this criterion and continued to do so for all subsequent visits, resulting in *N*_tr_ = 115 youth. The *T*_0,*i*_ for this example is thus the visit age at which this was initially reported for the *i*th participant. The control group consists of *N*_co_ = 513 adolescents who have never reported more than 12 binges in any year. Note, individuals were dropped who 1) had a brain anomaly in their MRI scan, or 2) had at least one visit reporting at least 12 binge drinking episodes in the past year but had a subsequent visit where they dropped below this threshold, resulting in a total sample size for the analyses of *N* = 628.

The baseline characteristics of the two groups are summarized in Table 2. The demographic characteristics were of comparable baseline ages across both groups, but there were a higher percentage of male and White individuals in the exposed group.

**Table 2:** Baseline demographic characteristics by group.

|  | Control | Exposed |
| --- | --- | --- |
| N | 513 | 115 |
| Baseline age in years (sd) | 15.79 (2.37) | 14.78 (1.72) |
| Female, % (n) | 52.6% (270) | 43.5% (50) |
| Asian, % (n) | 7.8% (40) | 7.8% (9) |
| Black, % (n) | 15.2% (78) | 5.2% (6) |
| Other race, % (n) | 5.3% (27) | 7.0% (8) |
| White, % (n) | 71.7% (368) | 80.0% (92) |

**Table 3:** Linear mixed-effects model for **standardized (z-scored)** superior frontal gray volume. Fixed effects estimate the association of heavy binge drinking (12+ episodes/year) with the standardized outcome while adjusting for standardized baseline intracranial volume, sex, visit, baseline age, and race. The model includes subject-specific random intercepts and random slopes for visit (`random = ∼visit | id`). Race coefficients are relative to the omitted reference race category.

| Predictor | Estimate | Std. Error | *p*-value |
| --- | --- | --- | --- |
| (Intercept) | 1.842 | 0.198 | < 0.001 |
| Heavy binge drinking (12+ episodes/year) | −0.073 | 0.020 | 0.00033 |
| Baseline intracranial volume (standardized) | 0.679 | 0.033 | < 0.001 |
| Male (vs female) | 0.130 | 0.062 | 0.035 |
| Visit (year) | −0.121 | 0.003 | < 0.001 |
| Baseline age (years) | −0.117 | 0.011 | < 0.001 |
| Race: African-American/Black | 0.345 | 0.125 | 0.0060 |
| Race: Asian | 0.246 | 0.138 | 0.0747 |
| Race: Caucasian/White | 0.240 | 0.109 | 0.0278 |

**Table 4:** Average treatment effect on the treated (ATT) over time with 95% confidence intervals.

| Time | ATT | 95% CI | $N$ |
| --- | --- | --- | --- |
| -3 | 38.1 | $[-5.1, 82.5]$ | 75 |
| -2 | -6.3 | $[-53.2, 40.6]$ | 71 |
| -1 | 11.9 | $[-34.6, 64.3]$ | 50 |
| 0 | -17.6 | $[-61.4, 25.3]$ | 139 |
| 1 | -47.7 | $[-109.4, 10.1]$ | 82 |
| 2 | -135.8 | $[-212.5, -58.0]$ | 49 |
| 3 | -259.2 | $[-411.3, -118.1]$ | 15 |

In prior work, we found associations between binge drinking in the past year and longitudinal gray matter volumes, with some of the strongest associations reported in the superior frontal region [10]. However, despite efforts to control for confounders, it remains possible that prior differences in gray matter volume trajectories at least partially account for the observed associations in that study. Thus, here we focus on the outcome of superior frontal gray matter volume(standardized) across up to 10 waves of annual follow-up.

We model each individual’s trajectory of standardized superior frontal gray volume (measured by magnetic resonance imaging at up to 9 annual visits) with our functional GSC approach to study the effect of heavy binge drinking(12+ episode a year), controlling for subject’s intracranial volume, sex, and race. The treated group had between 1 and 9 visits, with a mean (median) of 6.07 (6) visits. The control group also had between 1 and 9 visits, with a mean (median) of 5.56 (6). The distribution of ages of onset for 12+ binge drinks in the treated group is shown in Figure 4.

**Figure 4.**
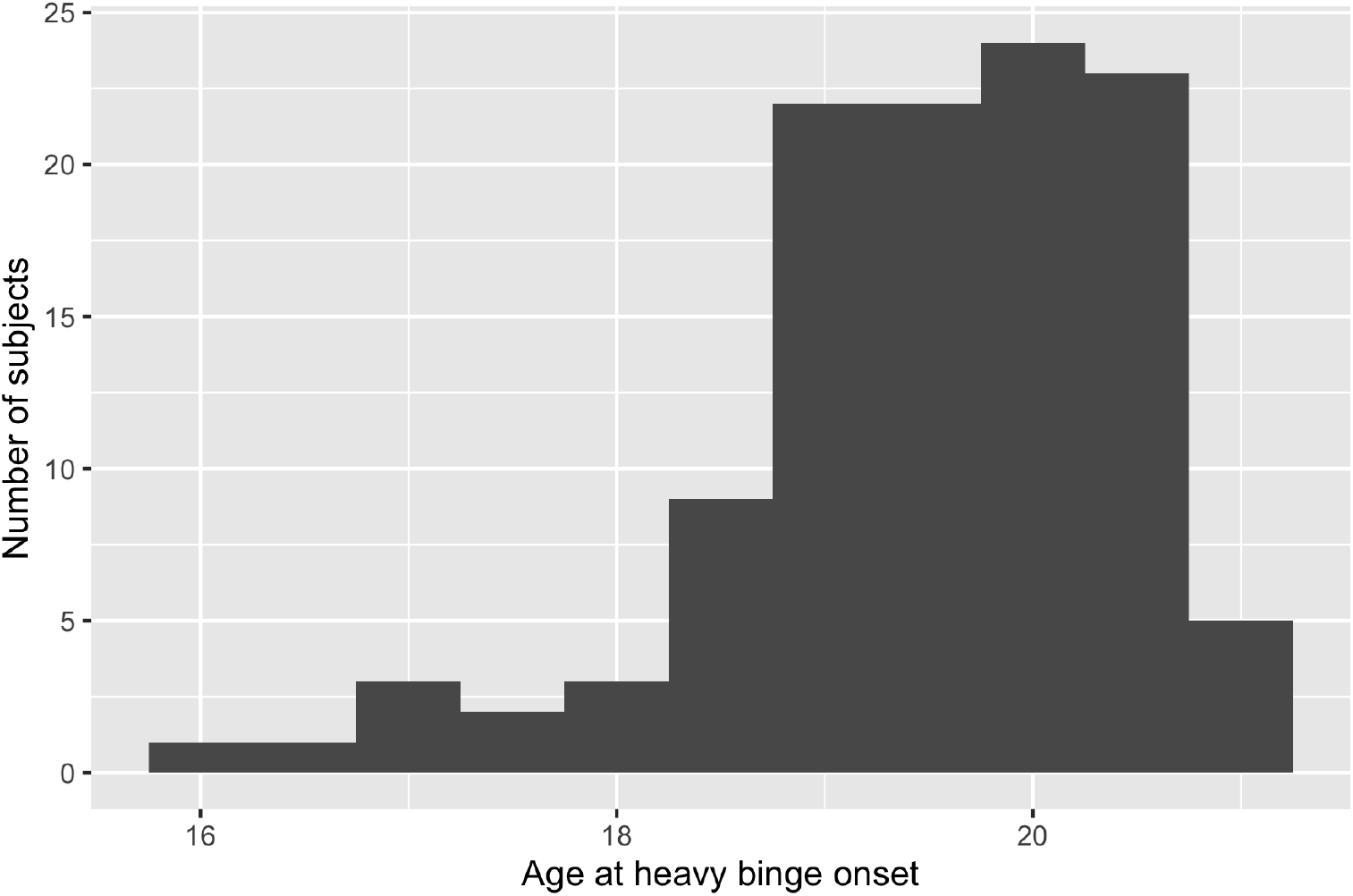
Age distribution (in years) of onset of reported heavy binge drinking behaviors in the *N* = 115 exposed participants.

**Figure 5.**
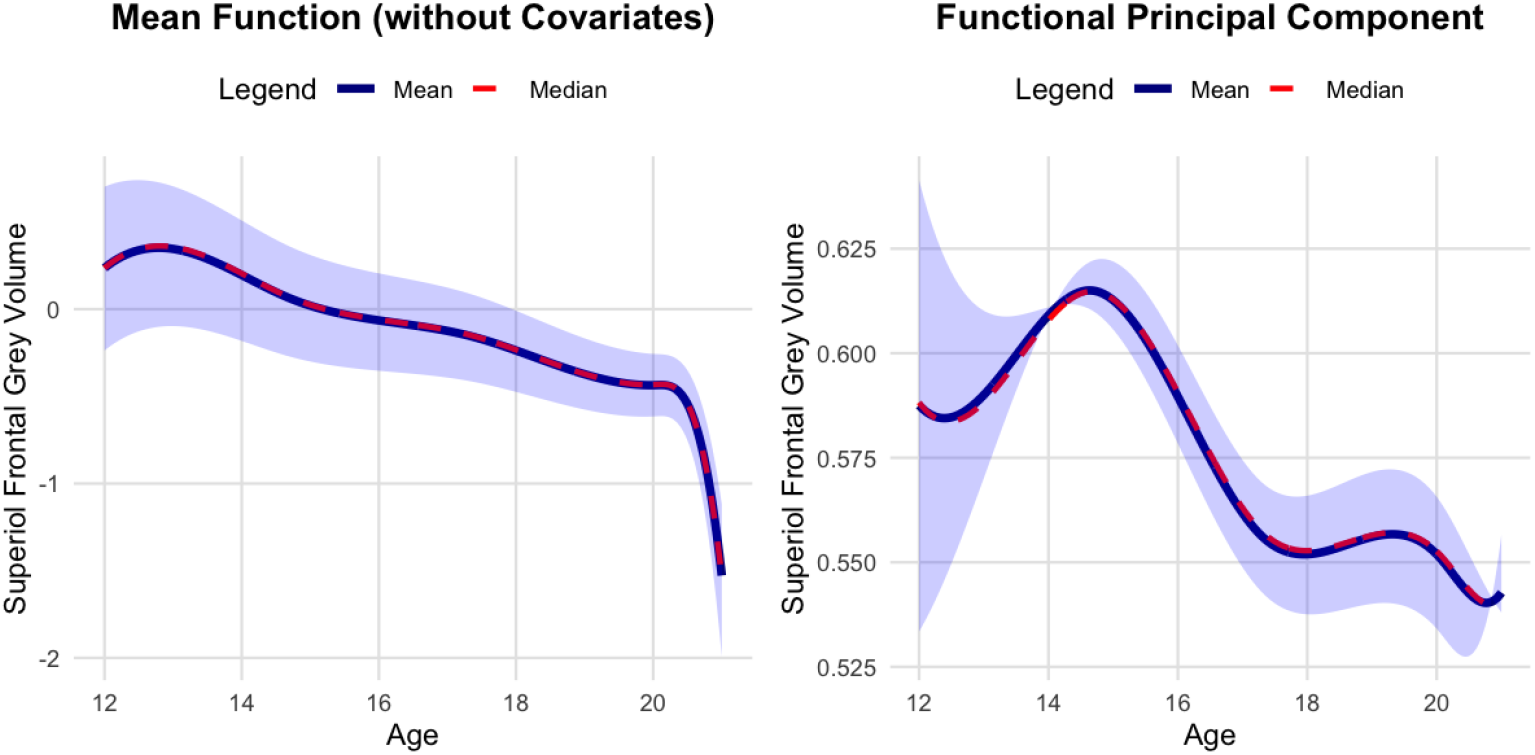
Estimated overall mean function 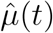 for superior frontal gray volume(standardized) (solid line) and the first functional principal component *f*_1_(*t*) (dashed line), from the control/pre-treatment data. The mean function suggests a decreasing trend with age. The first FPC represents an additional deviation pattern (individuals differ by how much they follow this pattern, scaled by their score *α*_*i*1_).

For exposed individuals, *T*_0,*i*_ is the year they first reported 12 or more binge episodes in the past year; before that year they are considered unexposed. We included as time invariant covariates: (1)baseline intracranial volume and (2) sex (3) race. These covariates help control for confounders and risk factors that could affect brain volume. We did not include any time-varying covariates. The FPCA was performed on the control group data, to ensure the latent factors represent unexposed developmental trajectories.

Figure 7 shows the PSIS-LOO (Pareto Smoothed Importance Sampling Leave-One-Out) [16][15] diagnostic for model fit for the final (one-FPC) specification. The resulting LOO estimates were elpd * loo = − 281.9 (SE = 90.8), LOOIC = 563.8 (SE = 181.5), with an effective number of parameters *p \** loo = 92.5 (SE = 10.4). The Pareto-*k* diagnostics indicate that most observations are well-behaved: 615/628 (97.9%) fall in the “good” range (*k* ≤ 0.63), 11/628 (1.8%) are in the “bad” range (0.63 *< k* ≤ 1), and only 2/628 (0.3%) exceed *k >* 1 (visible above the upper reference line in Figure 7), suggesting a small number of influential observations but no widespread instability. Consistent with the model-comparison results (the second FPC did not yield a statistically meaningful improvement in out-of-sample predictive performance), we retained a single FPC for parsimony.

Using the fitted model, we estimated the ATT on standardized (z-scored) superior frontal volume associated with heavy binge drinking. Figure 6(A) plots the estimated ATT as a function of time relative to the onset of heavy binge drinking. Time 0 on the x-axis corresponds to the measurement just before an individual crossed the threshold of 12 binges/year, and the post-exposure period corresponds to follow-up years floored to the nearest year after that point. The solid line is the posterior mean ATT and the shaded band is the 95% posterior predictive interval. We see that in the pre-exposure period, the ATT estimates are close to zero and their intervals include zero, indicating no detectable differences between exposed and control groups prior to heavy binge drinking. At Time 0, the estimated ATT is also not statistically distinguishable from zero (ATT ≈ −0.010; 95% PI: [−0.036,; 0.015]; original (non-standardized) units: −17.65, [−61.44,; 25.34]). In the first year after heavy binge drinking is reported, the estimated effect remains non-significant (ATT ≈ −0.028; 95% PI: [−0.063,; 0.006]; original (non-standardized) units: −47.66, [−109.40,; 10.14]). By the subsequent year, heavy binge drinking (12+ episodes in the past year) is associated with a mean decrease in standardized superior frontal volume of about −0.079 (95% PI: [−0.123,; −0.034]; original (non-standardized) units: −135.81, [−212.52,; −57.97]) compared to the counterfactual of no heavy binge drinking. Three consecutive years of heavy binge drinking are associated with an average standardized volume loss of about −0.150 (95% PI: [−0.238,; −0.068]; original (non-standardized) units: −259.16, [−411.33,; −118.15]) relative to the no-binge counterfactual. These intervals do not overlap zero, suggesting a statistically significant detrimental effect of sustained binge drinking on this brain structure.

**Figure 6.**
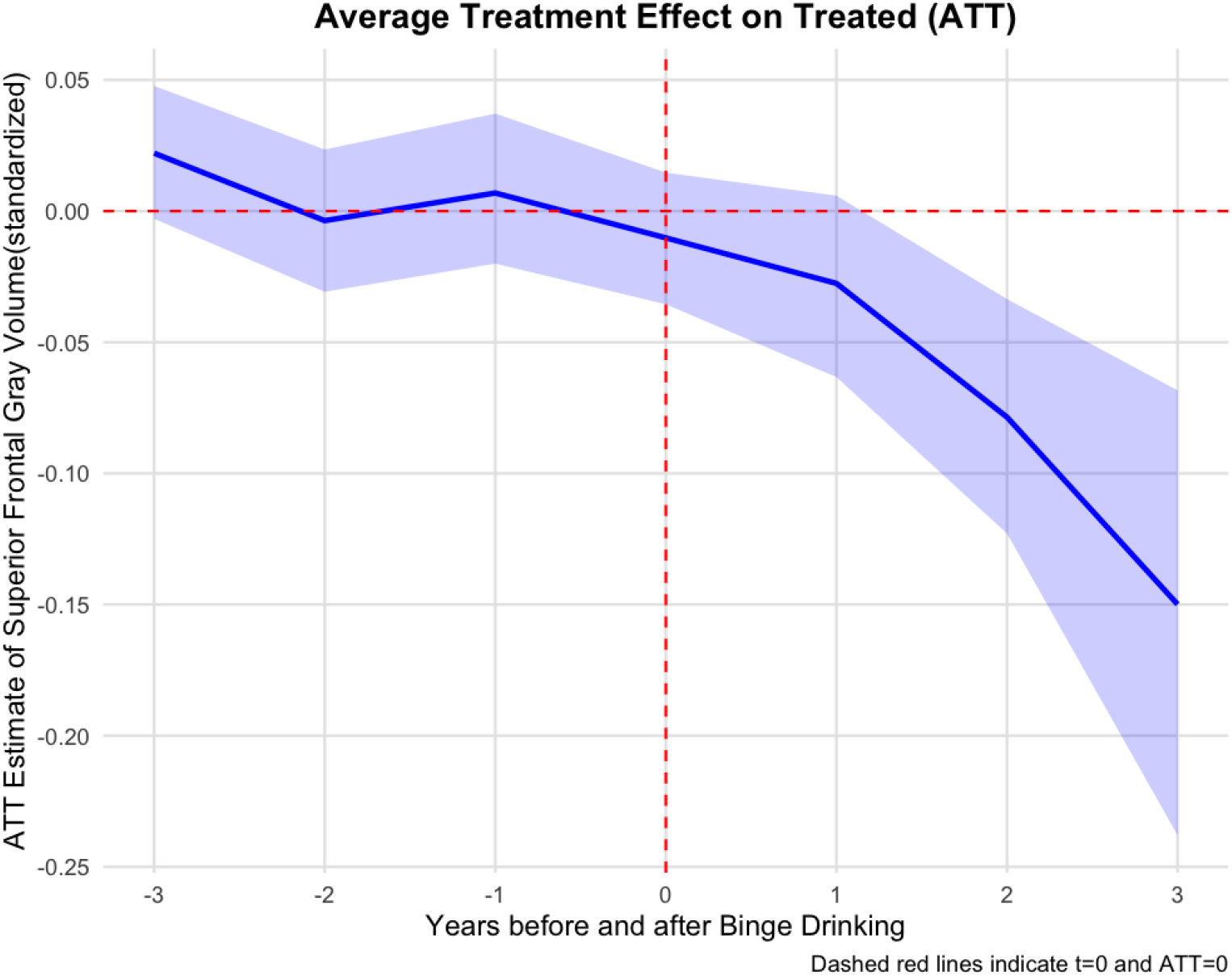
Estimated average treatment effect on the treated (ATT) for superior frontal gray volume(standardized) as a function of time relative to onset of heavy binge drinking. Time 0 is just before exposure (crossover to 12+ binges/year). Shaded region is the 95% predictive interval. A drop in volume is observed following exposure.

**Figure 7.**
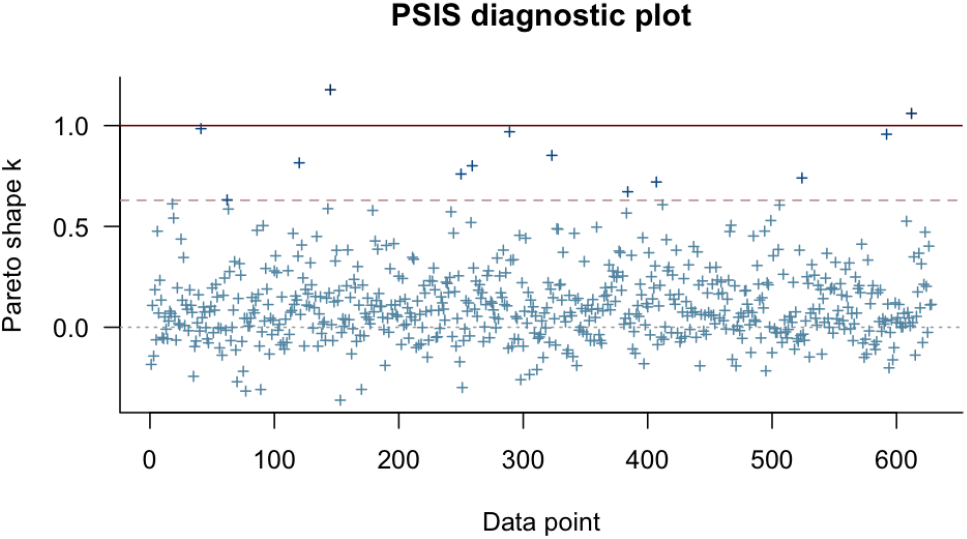
Model diagnostics: Pareto *k* values from PSIS-LOO for the one-FPC model fit. Most *k* values are well below 0.7 (dotted line), indicating a reliable fit. Only a few subjects have slightly higher *k*, suggesting minor outliers but no major lack of fit.

We also visualized the individual fit by randomly plotting 16 subjects fitted counterfactual and observed outcome(Figure 9). Across the 16 subjects, the fitted counterfactual curves generally show a gradual decline in superior frontal gray matter volume(standardized) with increasing age. Before the red vertical line (age of first heavy binge drinking exposure), the observed values (dots) typically track the counterfactual trend and fall within the 95% prediction interval. After the exposure point, many subjects’ observed volumes sit below the counterfactual curve, indicating lower-than-expected volume relative to what the pre-exposure trajectory would predict, consistent with a negative ATT estimate.

We also fit the time-varying coefficients for baseline covariates in the model (standardized baseline intracranial volume, sex, and race). We plotted the effects of baseline age and standardized baseline intracranial volume in Figure 8. Across ages, baseline ICV shows a consistently positive and statistically significant association with standardized superior frontal grey volume: the estimated coefficient remains above zero throughout the age range, and the 95% prediction interval does not overlap zero, indicating robust evidence that larger ICV is associated with higher standardized superior frontal grey volume at each age. In contrast, the sex effect (female relative to male) is not consistently statistically significant across development. Although the point estimates trend negative through mid–late adolescence, the 95% prediction interval overlaps zero for most (and likely all) ages, suggesting limited evidence for a reliable difference in superior frontal grey volume(standardized) by sex once other covariates are accounted for. The race covariates are not statistically significant as well (their 95% prediction intervals overlap zero), and the corresponding time-varying coefficient trajectories are visualized in the Appendix.

**Figure 8.**
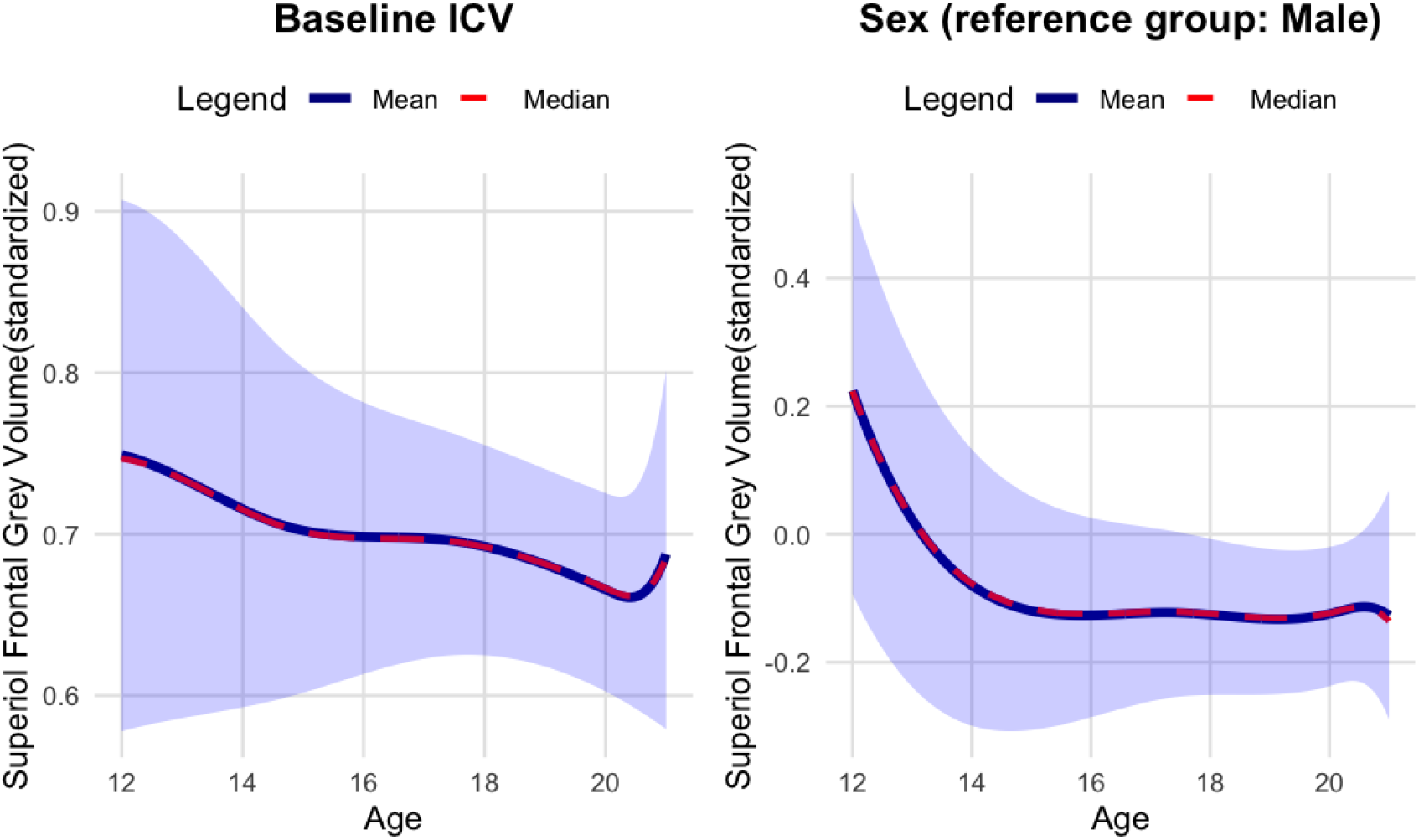
Age-varying covariate effects on standardized superior frontal grey volume. The left panel shows the estimated coefficient relating baseline intracranial volume (ICV) to superior frontal grey volume(standardized), and the right panel shows the estimated coefficient for sex (reference group: male). Solid blue lines denote the posterior mean coefficient and dashed red lines denote the posterior median across age; the shaded band indicates the 95% uncertainty interval.

**Figure 9.**
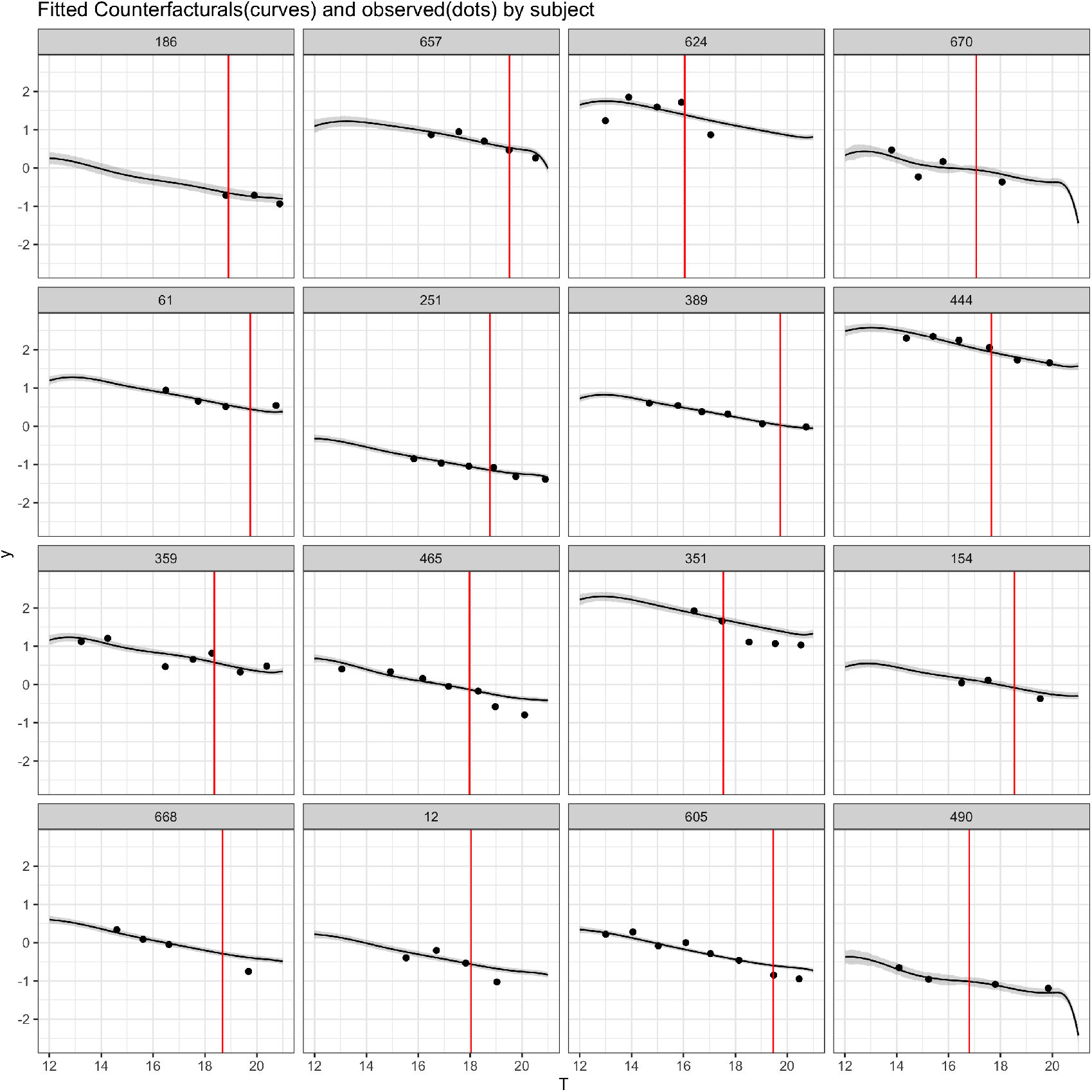
The individual fit of randomly selected 16 subject’s counterfactual fit(lines) and observed value(dots). Shaded area is the 95% prediction interval. The red vertical line indicates the age first exposed to heavy binge drinking.

**Figure 10.**
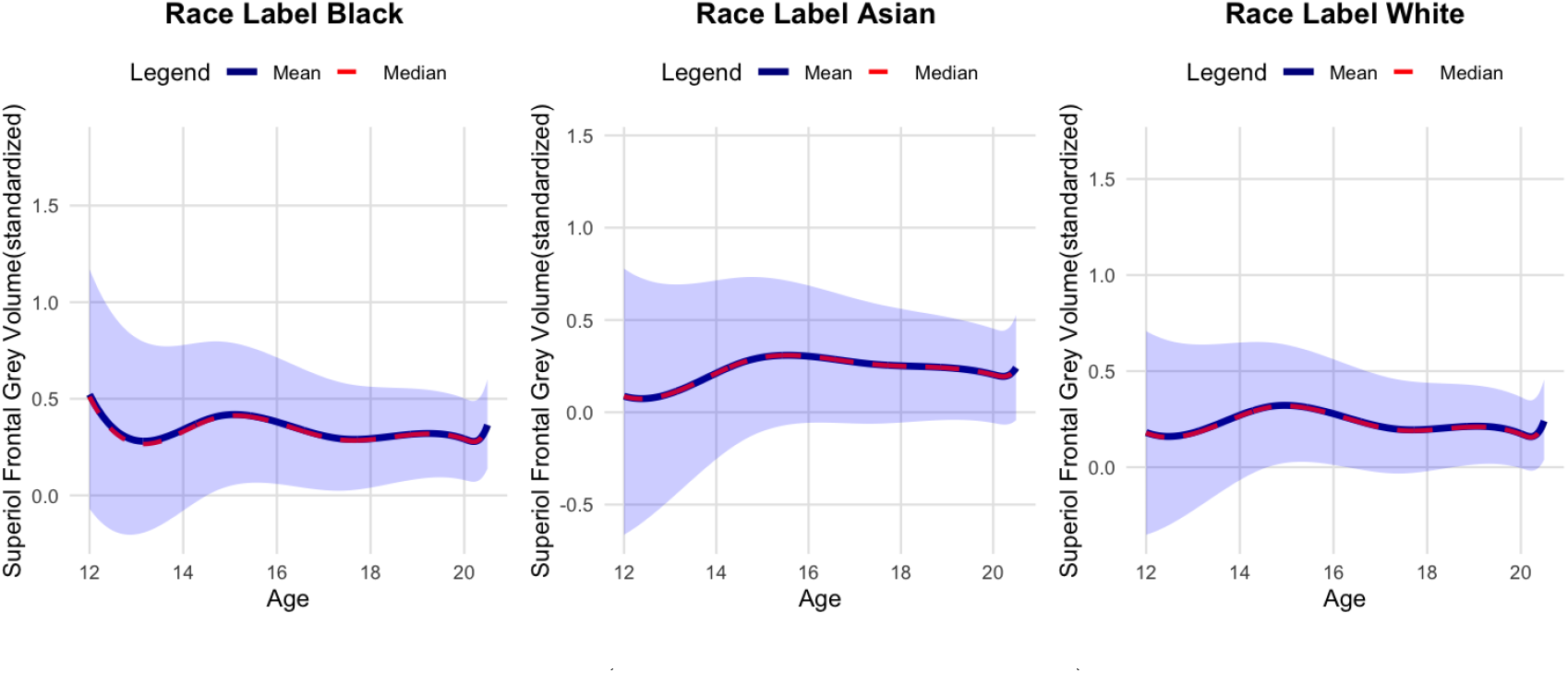
Age-varying racial(reference group: other) effects on standardized superior frontal grey volume. Solid blue lines denote the posterior mean coefficient and dashed red lines denote the posterior median across age; the shaded band indicates the 95% uncertainty interval.

To benchmark our functional GSC estimates against a more conventional longitudinal approach, we also fit a linear mixed-effects model (LME) for the standardized (z-scored) superior frontal gray volume, including heavy binge drinking (12+ episodes/year) and adjusting for baseline ICV, sex, visit, baseline age, and race, with subject-specific random intercepts and random slopes for visit. In this LME, heavy binge drinking was associated with a statistically significant average decrease of −0.073 SD (SE = 0.020, *p* = 3.3 × 10^−4^), which implicitly assumes a single, constant effect of binge drinking across follow-up time. In contrast, our functional GSC/event-time ATT analysis estimates a time-varying treatment effect relative to the onset of heavy binge drinking, using the pre-exposure period to anchor the counterfactual trajectory. Consistent with the identifying assumptions, the pre-exposure ATT estimates are near zero and not statistically significant (e.g., Time −3: 0.022 SD, 95% PI [−0.003,; 0.048]; Time −2: −0.004 SD, [−0.031,; 0.023]; Time −1: 0.007 SD, [−0.020,; 0.037]), and there is no detectable difference at Time 0 (−0.010 SD, [−0.036,; 0.015]). Post-onset, the effect increases in magnitude with continued exposure, consistent with a cumulative detriment: at 1 year the ATT is not significant (−0.028 SD, [−0.063,; 0.006]), but by 2 years it is significantly negative (−0.079 SD, [−0.123,; −0.034]), and by 3 years it is larger still (−0.150 SD, [−0.238,; −0.068]). Taken together, the LME’s single binge coefficient summarizes an average decrement (roughly comparable in magnitude to the 2-year ATT), whereas our method shows that the association is not immediate and constant, but instead appears to accumulate over up to three years of sustained heavy binge drinking.

In summary, our method finds evidence that heavy binge drinking in late adolescence is associated with a significant reduction in superior frontal gray matter volume, even after controlling for sex and overall brain size. The continuous-time functional approach allowed us to utilize all available data points and capture individual developmental trajectories. By comparing each binge-drinking youth to a data-adaptive synthetic trajectory (learned from controls), we obtained an estimate of the counterfactual path and thus the causal effect. These findings underscore the potential neurotoxic effects of sustained heavy alcohol use during a critical period of brain development.

## 5 Discussion

In this paper, we developed a generalized synthetic control algorithm tailored for sparse functional data. By incorporating Functional PCA into the synthetic control framework, our method can flexibly model irregularly sampled trajectories and estimate causal effects without requiring strict alignment of time points across individuals. Through simulations, we demonstrated that the approach performs well in recovering treatment effects under a variety of scenarios, and provides well-calibrated uncertainty intervals. The application to neuroimaging data illustrated how the method can leverage information from all available time points to detect a plausible causal impact (here, the effect of binge drinking on brain volume) that might be missed or attenuated by methods that aggregate or ignore timing information.

Our approach inherits some limitations common to both synthetic control and functional data models. In particular, it relies on the assumption that the control group (in combination with the latent factors) can adequately reconstruct the treated units’ counterfactual trajectories. If there are unmeasured confounders or latent patterns that are unique to the treated group and not present in the controls, our estimates could still be biased. We partially mitigate this by including relevant covariates and using a flexible latent basis, but unobserved group-specific shocks remain an identifying concern (as with any non-randomized comparison). Another limitation is the need to choose the number of FPCs *k* and the spline basis dimension *q*; we used LOOIC to guide the choice of number of FPC’s, but model misspecification in these could affect results. In practice, one should examine whether results are sensitive to including an extra FPC or changing the basis flexibility.

There are several avenues for extension. One could incorporate nonlinear or time-varying treatment effects by allowing *δ*(*t*) to be subject-specific or dependent on observed moderators (e.g, age of onset), moving toward a personalized treatment effect function. The framework can also be extended to multiple outcomes (multivariate functional data) by using multi-dimensional FPCA. Additionally, one could consider alternative prior structures or penalization for the functional components to improve robustness (e.g., sparsity-inducing priors on Θ if one expects only a few basis functions to matter). Another promising direction is to integrate recent advances in functional data warping, in case individuals’ trajectories are similar in shape but misaligned in time (though in our setting alignment is defined by real time and an intervention point, so this was less of an issue).

Despite these caveats, we believe the proposed methodology fills an important gap for causal inference in longitudinal studies with irregular observation patterns. It includes the strengths of synthetic controls (robust causal effect estimation under the weak trends assumptions) with those of functional data analysis (flexible modeling of irregularly-sampled curves). This makes it well-suited for biomedical applications and other scenarios where follow-up schedules vary by subject or data collection is sparse. The use of Bayesian estimation provides a convenient way to quantify uncertainty in complex models. We provide software implementing this method at https://github.com/l66shao/GSC-FPCA, so that researchers can apply it to their own data. As increasingly rich longitudinal datasets become available (often with uneven measurement times), tools that can exploit the full temporal information will be crucial for drawing credible causal conclusions.

## AcknowleWordgement

The work was partly supported by the National Institute of Health (DA057567, AA021697, MH128959) and the 2024 Stanford HAI Hoffman-Yee Grant. Collection and curation of the NCANDA data was supported by the U.S. National Institute on Alcohol Abuse and Alcoholism with co-funding from the National Institute on Drug Abuse, the National Institute of Mental Health, and the National Institute of Child Health and Human Development (Grant Numbers AA021697, AA021695, AA021692, AA021696, AA021681, AA021690, AA021691).

## Appendix

### 5.1 Appendix A: Gibbs Sampler Derivation

For completeness, we include the full conditional posterior derivations for the Gibbs sampling steps described in Section 3.2. Using the notation from that section, recall 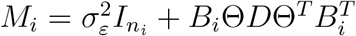.

Combining the likelihood from the model (9) with the priors above, we derive a Gibbs sampler to obtain posterior samples of the parameters and latent variables. Due to the conjugate structure of the model, most conditional posterior distributions take a convenient form. We briefly outline the forms of these full conditional distributions here (see Appendix for detailed derivations).

Let 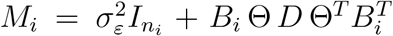 denote the marginal covariance of *Y*_*i*_ (after integrating out *α*_*i*_). The conditional posterior for the mean function coefficients *θ*_*µ*_ is multivariate Gaussian:

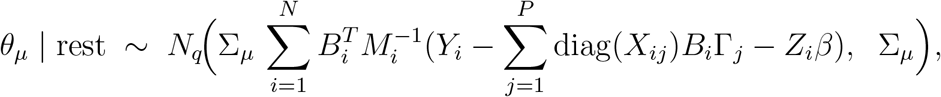

where 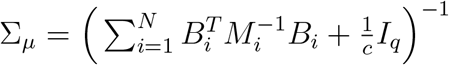.

For each subject’s FPC scores *α*_*i*_, the conditional posterior is:

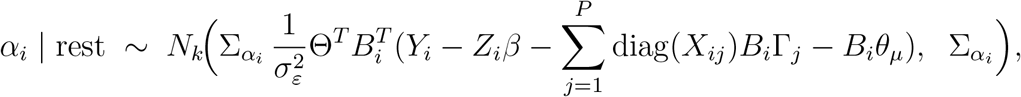

where 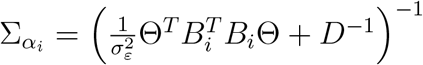.

For each FPC basis coefficient vector Θ_·*j*_ (the *j*th column of Θ), the conditional posterior (enforcing orthogonality via a Gram-Schmidt step) is:

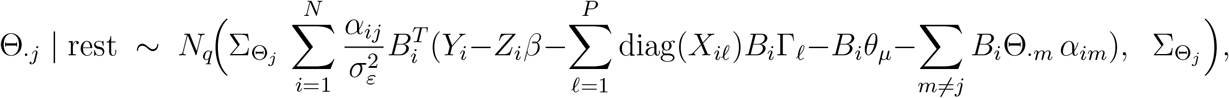

with 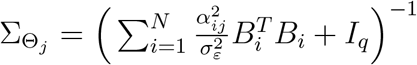.

The residual variance 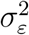 has an inverse-gamma conditional posterior:

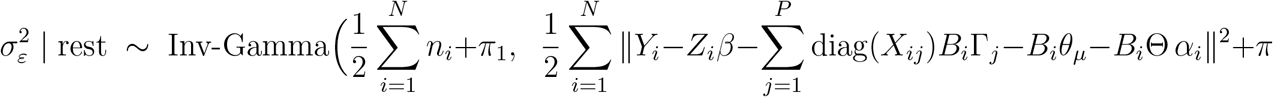

where ∥ · ∥^2^ denotes squared Euclidean norm.

For each latent factor variance *d*_*j*_ (diagonal element of *D*), the conditional posterior is:

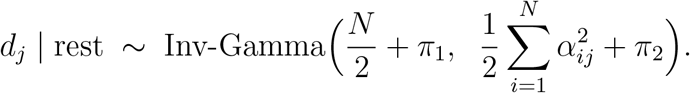

Finally, for each time-varying covariate coefficient Γ_*j*_, the conditional posterior is multivariate normal:

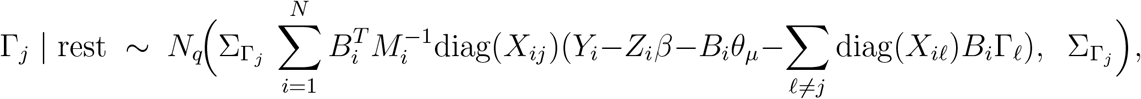

where 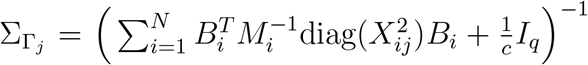 (here 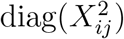) indicates the diagonal matrix of squared covariate values, since in the Gaussian linear model the effective design for Γ_*j*_(*t*) is weighted by *X*_*ij*_(*t*) at each time).

We iterate through these conditional draws in a Gibbs sampler. The orthonormality constraint on Θ is enforced by re-orthonormalizing the columns of Θ after each update (this step is of negligible computational cost given the low dimension *q* of the basis). After a burn-in period, we obtain posterior samples for all parameters and latent variables. From these, we can compute the posterior predictive distribution of each treated unit’s counterfactual 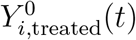 in the post-treatment period, and hence the posterior of the treatment effect *δ*_*i*_(*t*). In practice, we report the posterior mean estimates and credible intervals for the ATT(*t*) at various time points.

#### Conditional posterior for *θ*_*µ*_

The conditional density for *θ*_*µ*_ (up to proportionality) is

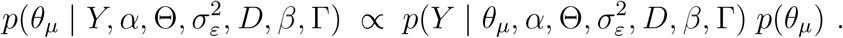

Combining the Gaussian likelihood and prior yields:

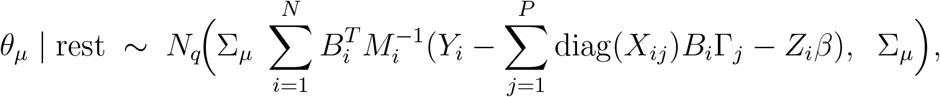

with 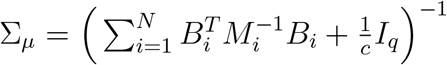.

#### Conditional posterior for *α*_*i*_

For each *i*, treating *α*_*i*_ as missing data in a linear mixed model, the complete-data likelihood is Gaussian in *α*_*i*_ and the prior for *α*_*i*_ is *N* (0, *D*). The conditional posterior is:

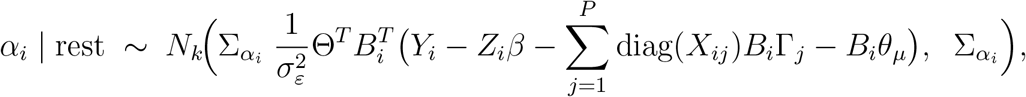

where 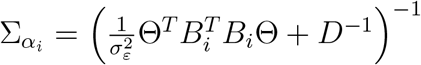.

#### Conditional posterior for Θ

Let Θ_·*j*_ denote the *j*th column of Θ (the coefficients for FPC *j*). Its conditional posterior (before enforcing orthonormality) comes from the likelihood of all *Y*_*i*_ depending on Θ_·*j*_ through *α*_*ij*_ and the *j*th component of *B*_*i*_Θ*α*_*i*_, combined with the *N* (0, *I*_*q*_) prior for Θ_·*j*_. Completing the square, we get:

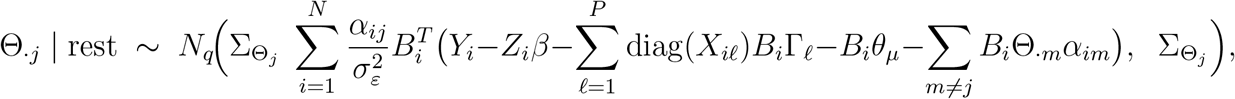

with 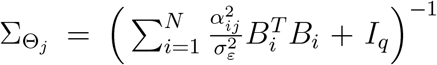. After sampling, we orthonormalize 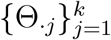 via a QR decomposition to ensure Θ^*T*^ Θ = *I*.

#### Conditional posterior for 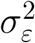

Treating all *α*_*i*_ as known (from the augmentation), the residual variance has an inverse-gamma posterior from the conjugate prior:

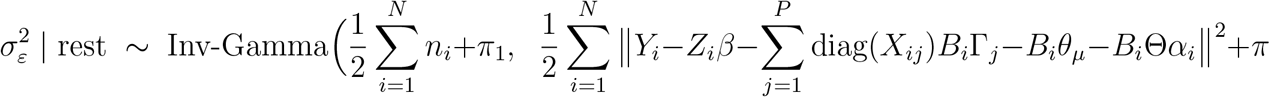

#### Conditional posterior for *d*_*j*_ (elements of *D*)

Each *d*_*j*_ (the variance of FPC score *α*_*ij*_) has a conjugate inverse-gamma posterior:

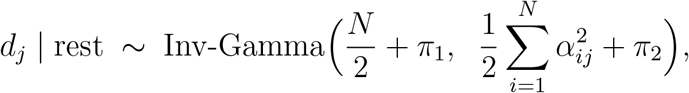

since *α*_*ij*_ ∼ *N* (0, *d*_*j*_) and prior *d*_*j*_ ∼ Inv-Gamma(*π*_1_, *π*_2_).

#### Conditional posterior for Γ_*j*_

The conditional posterior for the coefficient function Γ_*j*_(*t*) (represented by basis coefficients Γ_*j*_) is:

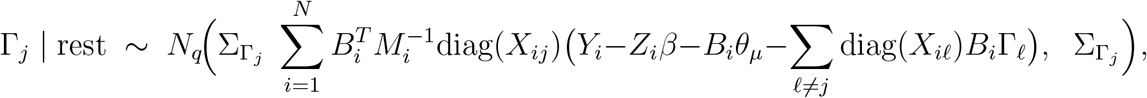

with 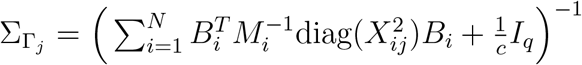.

### 5.2 Appendix B: Empirical Example

#### 5.2.1 Average Treatment Effect on Superior Frontal Gray Volume

#### 5.2.2 Coefficient Plot for Race

